# Quantification of subclonal selection in cancer from bulk sequencing data

**DOI:** 10.1101/096305

**Authors:** Marc J. Williams, Benjamin Werner, Christina Curtis, Chris P Barnes, Andrea Sottoriva, Trevor A Graham

**Affiliations:** Evolution and Cancer Laboratory, Barts Cancer Institute, Queen Mary University of London, London, UK.; Department of Cell and Developmental Biology, University College London, London, UK.; Centre for Mathematics and Physics in the Life Sciences and Experimental Biology (CoMPLEX), University College London, London, UK.; Centre for Evolution and Cancer, The Institute of Cancer Research, London, UK.; Departments of Medicine and Genetics, Stanford University School of Medicine, Stanford, CA 94305, USA; Stanford Cancer Institute, Stanford University School of Medicine, Stanford, CA 94305, USA

## Abstract

Recent studies have identified prevalent subclonal architectures within many cancer types. However, the temporal evolutionary dynamics that produce these subclonal architectures remain unknown. Here we measure evolutionary dynamics in primary human cancers using computational modelling of clonal selection applied to high throughput sequencing data. Our approach simultaneously determines the subclonal architecture of a tumour sample, and measures the mutation rate, the selective advantage, and the time of appearance of subclones. Simulations demonstrate the accuracy of the method, and revealed the degree to which evolutionary dynamics are recorded in the genome. Application of our method to high-depth sequencing data from gastric and lung cancers revealed that detectable subclones consistently emerged early during tumour growth and had considerably large fitness advantages (>20% growth advantage). Our quantitative platform provides new insight into the evolutionary history of cancers by facilitating the measurement of fundamental evolutionary parameters in individual patients.

## Introduction

Carcinogenesis is the result of a complex process of Darwinian selection for malignant phenotypes^1,2^. The evolutionary process is driven by the accumulation of genetic alterations that allow cells to evade normal homeostatic regulation and prosper in changing microenvironments. High throughput genomics has shown that tumours across all cancer types are highly heterogeneous^3^, to the point that each cell may potentially be genetically unique^4^, thus leading to complex clonal architectures within tumours^5^. However, because longitudinal observation of tumour growth remains impractical, the temporal evolutionary dynamics that produce those clonal architecture remain undetermined. Knowledge of these evolutionary dynamics is necessary to infer future evolutionary trajectories and modes of relapse.

Studying the temporal process of cancer evolution is challenging because molecular information is usually collected from an individual’s cancer at a single time point, typically at resection. However, the subclonal architecture of a cancer – as measured by the pattern of intra-tumour genetic heterogeneity (ITH) – is directly determined by the unobservable evolutionary dynamics. Thus, given a realistically constrained model of clonal expansion during tumour evolution, the pattern of ITH in a tumour can be used to infer the most probable evolutionary trajectory of that tumour. ITH represented within the distribution of variant allele frequencies (VAF), which is measured by high coverage sequencing of cancer samples, is particularly amenable to such an approach.

We have previously shown that under a neutral evolutionary model (e.g. in the absence of subclonal selection), the VAF distribution has a predictable form that is observed in ~30% of cases from multiple cancer types^6^. However, the majority of samples (~70% of cancers analysed) showed VAF distributions that were not consistent with neutral evolution.

Here, using a stochastic model of subclone evolution in cancer and Bayesian inference, we identify the signature of selection in the cancer genome and quantify the evolutionary dynamics of non-neutrally evolving tumours.

## Results

### Theoretical framework

We developed a stochastic simulation of tumour growth that accounts for subclonal selection (Figure 1 and methods). At each division, a cell divides to produce either 0 or 2 surviving offspring with predefined probabilities, and daughter cells acquire new mutations at rate *μ* mutations per cell per division (figure 1A). The fitness advantage of a mutant subclone is defined by the ratio of net growth rates between the fitter mutant (*λ_m_*) and the background host population (*λ_b_*)

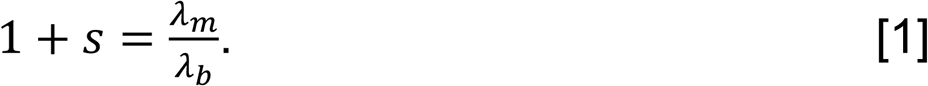

**Figure 1.**
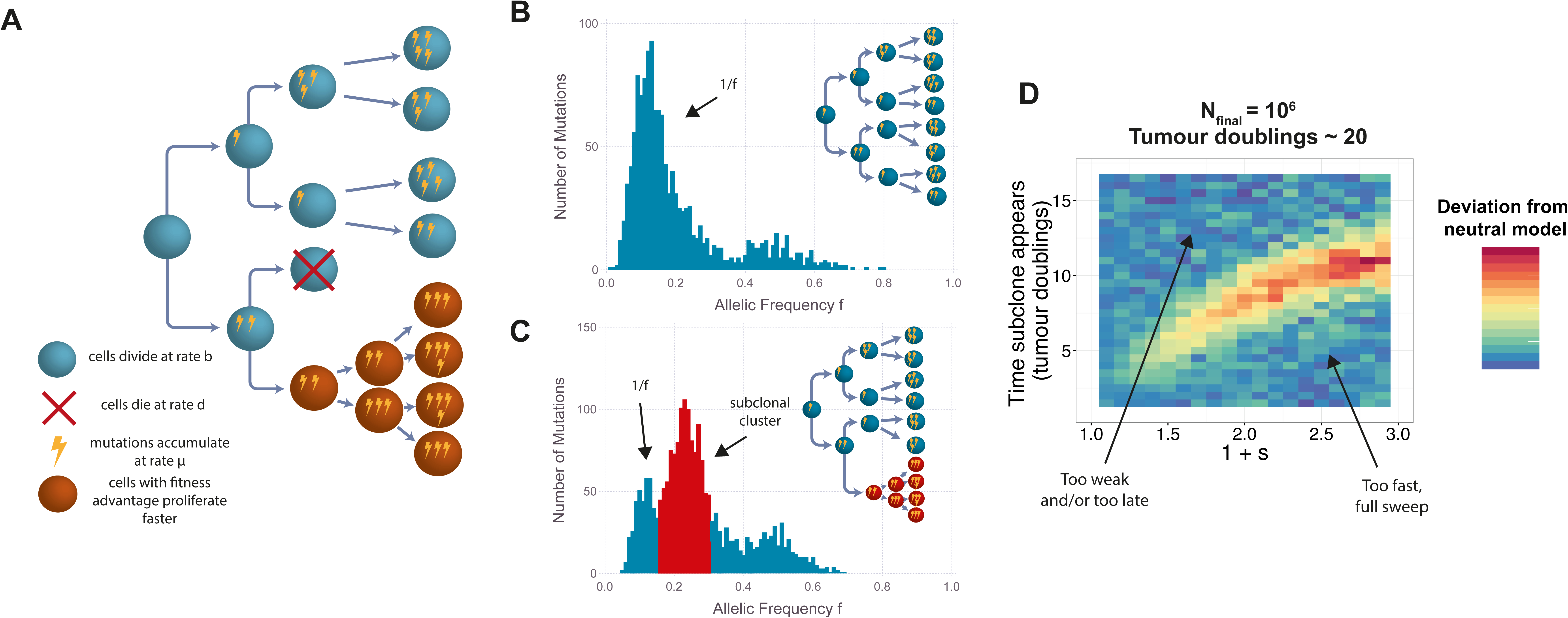
A We model tumour growth using a branching process where cells have a birth rate and a death rate, mutations accumulate as the tumour grows and cells with fitness advantages grow at a faster rate than the host population. The variant allele frequency is a consequence of how a tumour grows, simulating a tumour with subclonal selection results in an additional peak, **B** compared to the neutral case, **C**. Using a test to detect deviations from the neutral model, we introduced fitter mutations at different time points with varying selection coefficients and found that early/and or very fit subclones results in detectable deviations from the neutral model, **D**. Tumours were simulated with a final population size of 10^6^, each pixel represents the average value for the metric (area between curves, see methods) over 50 simulations.

This definition provides an intuitive interpretation for the fitness coefficient *s*, for example, an *s* of 1 implies that the mutant cell population grows twice as fast as the host tumour population. With the fitness coefficient *s* = 0, we have that *λ_m_*=*λ_b_* and the subclone evolves neutrally. Within the model, neutral evolution leads to a VAF distribution characterised by a 1/f-distributed subclonal tail of mutations^6^ (Figure 1B), whereas clonal selection produces characteristic ‘subclonal clusters’ within the VAF distribution that have been identified in previous analyses^7^ (figure 1C). Importantly, as neutral mutations continue to accumulate within each subclone, the 1/f-like tail is also present in tumours with selected subclones^8^ (figure 1C).

A mathematical analysis of the model indicates how subclonal clusters encode the underlying subclone evolutionary dynamics: the mean VAF of the cluster is a measure of the relative size of the subclone within the tumour, and the total number of mutations in the cluster indicates the subclone’s relative age. Together, these two measures allow the fitness advantage *s* to be estimated^9^.

We define t_0_=0 the time when the first transformed cancer cell begins to grow. At a later time t_1_, a cell in the tumour acquires a ‘driver’ somatic alteration that confers a fitness advantage, giving rise to a new subclone that expands faster than the other tumour cells. We note that the driver need not be a genetic mutation, but could be epigenetic or even microenvironmentally determined. The number of mutations acquired by the founder cell of the fitter subclone is

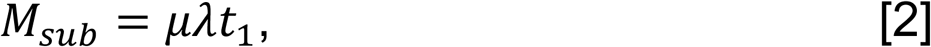

where *λ* = log (2) if we measure time in units of tumour volume doublings. We have previously shown that the effective mutation rate (the number of mutations for every newly generated lineage) can be estimated from the 1/f tail^6^. For a subclone that emerges at time t_1_ we would expect to observe *M_sub_* mutations at a frequency *f_sub_* which given some sequencing noise will present as a cluster of mutations with a mean *f_sub_* in the VAF distribution. Therefore, equation [2] allows us to estimate t_1_, the time when the subclone appeared.

N_sub_(t) and N_background_(t) represent the population size of the subclone and background populations at time *t*. The frequency of the subclone in the tumour at time *t_end_* when the tumour is resected is given by

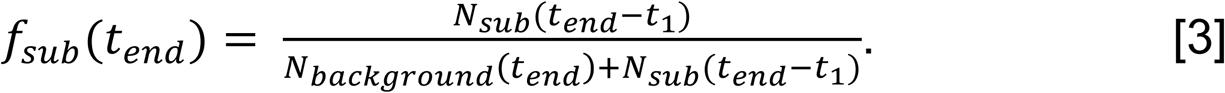

Under exponential tumour growth we have

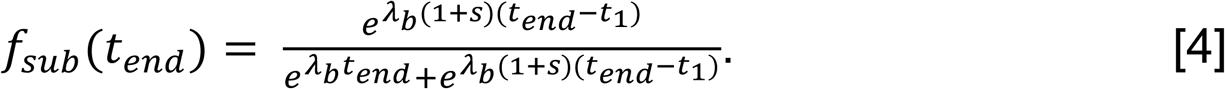

Solving for the fitness advantage *s* gives

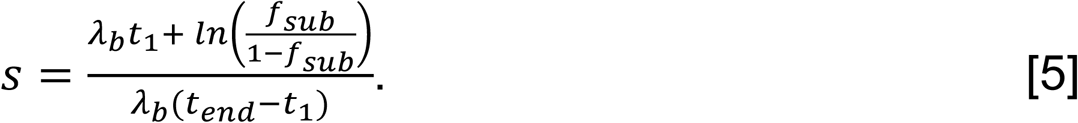

Therefore, given an estimate of the age of the tumour, t_end_ (for example assuming the final population size is 10^9^, we can calculate t_end_ via *2^t_end_^ =*(1 – *f_sub_*)×10^9^) then equations [2] and [5] provide a means to measure the selective advantage of a subclone directly from the VAF distribution (figure 2A). In the case where we have multiple subclones, equation [5] takes a slightly modified form (supplementary note).

**Figure 2.**
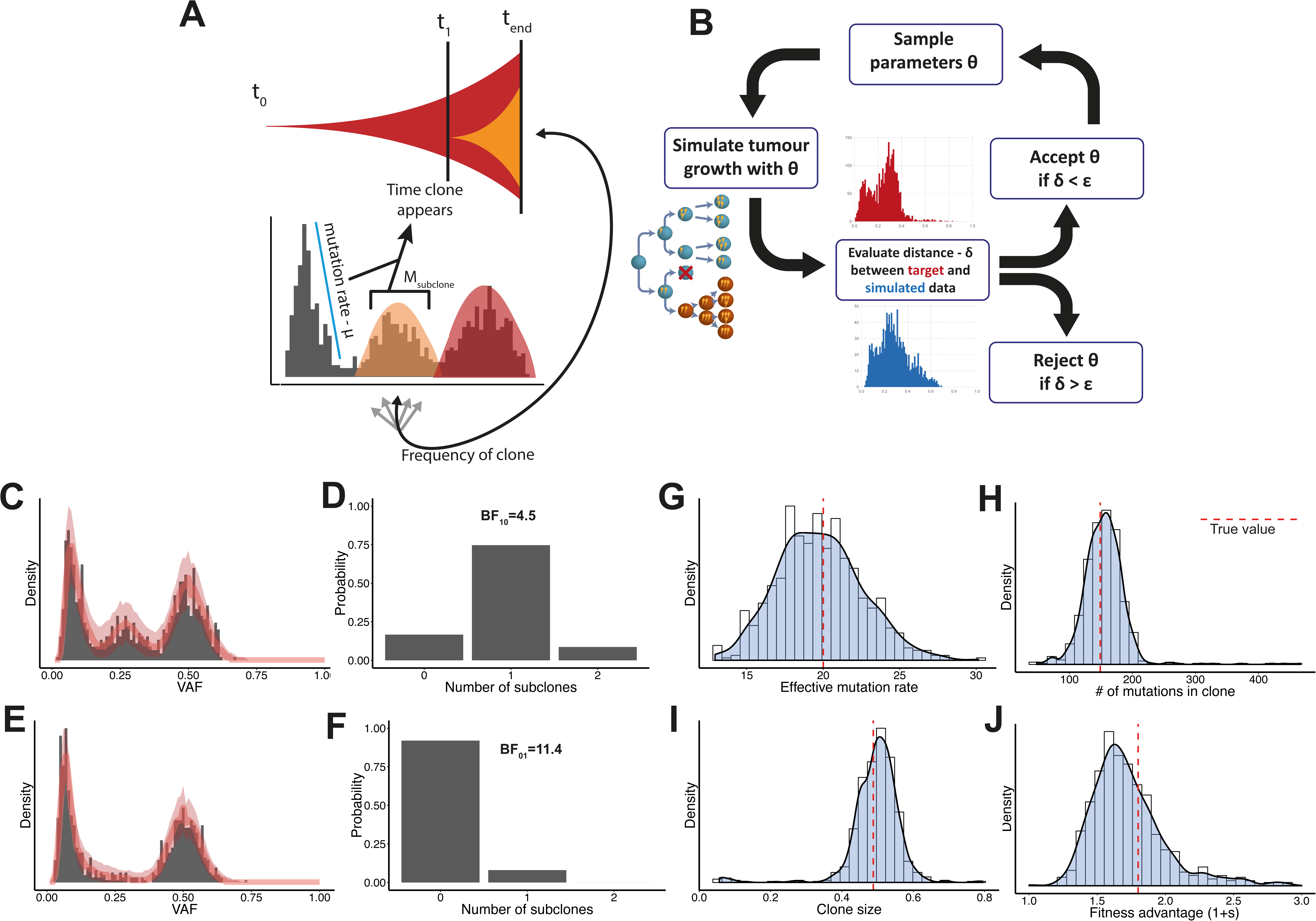
A Extracting the mutation rate, the number of mutations in the subclone and the frequency of the subclone from the VAF distribution allows us to first measure the age of a subclone which we then use to measure the selective advantage of a subclone. We used Bayesian statistical inference together with our simulation to measure these parameters, where we simulate our model many times with different parameters to find those parameters that produce synthetic datasets that closely match the target data, **B**. Applying our inference scheme to a simulated dataset with one subclone, **C** and a neutral simulated dataset **E** we were able to correctly identify the most probable number of subclones **D**, **F**. As well as accurately measure the effective mutation rate **G**, the number of mutations in the subclone **H**, the frequency of the subclone in the population I and its fitness advantage J. Simulation parameters: μ=5/division, β=0.25, time clone appears = 5.2 (tumour doubling times), number of mutations in clone = 149, 1+s=1.8 (b_H_=0.69, d_H_ = 0.52, b_F_ = 0.733, d_F_ = 0.42), frequency of subclone = 0.49, final population size = 105. Red line in panels **C** and **E** are the median histograms from the simulations that passed the ABC inference, shaded areas are the 95% intervals.

### Limits of detectability of subclonal selection

The ability to detect and quantify selection in human cancers naturally depends on the strength of the signal and the resolution of the data at hand. Because the population in tumours is expanding, the later a subclone appears, the fitter it has to be to grow to a detectable size before the tumour is removed from the body and studied in the lab^10^. Consequently, the combined effect of the fitness advantage of a subclone and the time of its appearance determine whether a clone will be of a detectable size. Moreover, in high throughput sequencing of cancer samples, the sequencing depth sets a lower limit on the size of observable subclonal mutations (e.g. ~5% for 100X depth sequencing^11^).

To determine how these evolutionary parameters and technical considerations constrain the ability to detect subclonal selection in the cancer genome, we developed a sensitive test that calculated the probability of observing a particular VAF distribution under neutral evolution. When the observed VAF distribution had a low probability of occurring under neutral evolution, we rejected the neutral model in favour of the alternative ‘subclonal selection’ hypothesis (see methods and supplementary figures 1,2). This analysis showed that only sufficiently early or very fit subclones are likely to be distinguishable from neutral evolution in moderate depth (100X) sequencing data (Figure 1D). In addition, the VAF distribution when the subclone is dominant (>90%) is indistinguishable from neutrality, as it is then the case that only neutral within-clone evolutionary dynamics are captured by the VAF distribution. In other words, once a fitter subclone has swept to near-fixation in a tumour, the tumour reverts to neutral dynamics.

### Accurate measurement of subclonal evolutionary dynamics in synthetic tumours

To infer evolutionary dynamics from VAF distributions, we implemented a Bayesian statistical inference framework (figure 2B & methods) that used our computational model of subclone evolutionary dynamics to simultaneously estimate all the parameters of interest from the sequencing data (principally the number of subclones, subclone fitness and time of occurrence, and the mutation rate). Importantly, this method allowed us to perform Bayesian model selection^12^ for the number of subclones within the tumour. This enabled us to calculate the probability that a given tumour contained 0 subclones (s=0, neutral evolution) or 1 or more subclones (non-neutral evolution).

In synthetic data (VAF distributions derived from computational simulations of tumour growth with known parameters), our framework accurately recovered the parameters governing tumour evolution in the presence of both subclonal selection (figure 2C,D) and neutral dynamics (figure 2E,F). In the case of subclonal selection, we were able to consistently recover the correct mutation rate (figure 2G), the number of mutations in the clone (figure 2H) and the size of the subclone (figure 2I), and via equation [5] we could infer an accurate posterior distribution for the fitness advantage of the subclone(figure 2J).

### Measuring subclonal selection in human cancers

We used our approach to quantify evolutionary dynamics in primary human cancers. We restricted our analysis to datasets were the depth of sequencing was very high to allow for accurate measurements of the clonal dynamics. To avoid the confounding effects of copy number changes, we exploited the hitchhiking principle^13^ and restricted our analysis to consider only single nucleotide variants (SNVs) that were located within diploid regions (see methods). Thus, upon adjusting for purity, we would expect to observe a ‘clonal cluster’ at vAF=0.5, and a potentially complex distribution of mutations with VAF<0.5 representing the subclonal architecture.

First, we applied our model to a high depth exome sequenced (>200X) lung adenocarcinoma dataset^14^. We used patient 4990 which had 5 samples from different sites sequenced. Sample 12 appeared to have a subclonal population (figure 3A). Bayesian inference found strong evidence in favour of one subclone, thus rejecting neutral evolution (figure 3A, Bayes Factor=9.1) and measured a median relative fitness of ~1.3 (95% credible interval:1.18-1.47) for the subclone over the background tumour population (figure 3B). In 4 other samples from patient 4990, our model identified neutral evolution as the most likely model (see supplementary figure 3). Sample 12 appeared to have copy number alterations on chromosome 3 that were not apparent in the other samples, suggestive that a copy number alteration may have driven the subclonal expansion (supplementary figure 4).

**Figure 3.**
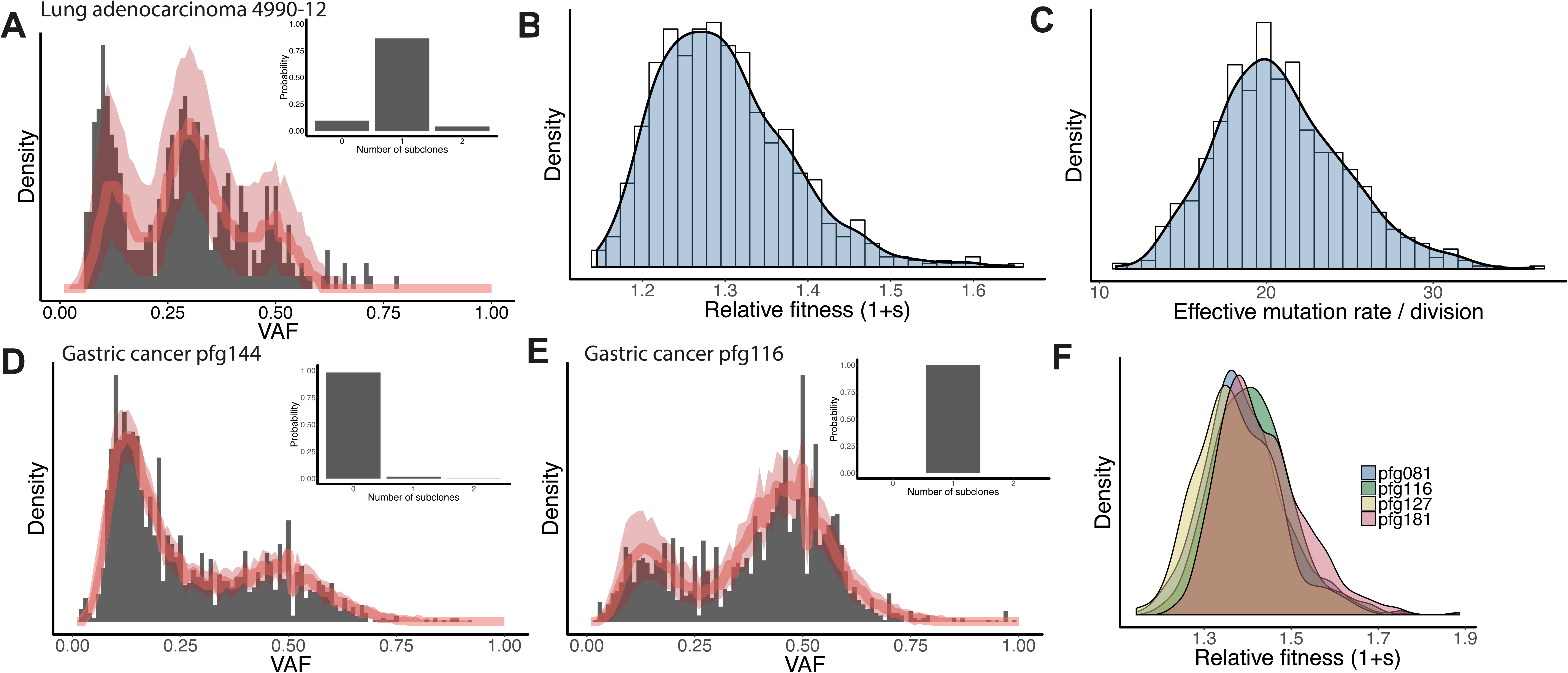
One sample from a lung adenocarcinoma dataset appeared to have a subclonal cluster **A**, our inference scheme identified a model with 1 clone as the most probably with a Bayes Factor of 9.1 in favour of this model over the neutral model. We inferred a median fitness advantage of 1+s~1.3, that is the clone grows 30% faster than the host tumour population **C**, and an effective mutation rate of 20/division/exome **C**. We found the stomach cancer sample pf144 to be consistent with a neutral model, **D** and sample pfg116 to be consistent with 1 subclone, although the subclone appears to be obscured by the clonal cluster **E**. Across 4 stomach cancer samples that showed evidence of a single subclone we observed similar fitness values **F**.

Next we applied the model to a whole genome sequenced gastric cancer dataset^15^. We applied the analysis to 10 samples that had high cellularity (>50%) and after removal of non-diploid regions contained a large number of mutations (>1000). Six of the samples showed strong evidence in favour of the neutral model (figure 3D and supplementary figure 5), while 4 samples had evidence of a subclone under differential selection (figure 3E and supplementary figure 5). As in the lung adenocarcinoma sample, we measured the relative fitness of the subclones to be >1.2 (20% advantage) in all 4 cases (figure 3F), and to have emerged early during tumour growth (supplementary figure 6). In these cases, there is no obvious subclonal cluster, possibly due to the comparably lower depth (~80X). Also we observed that the clonal cluster often appear to have more mass on the left hand side (supplementary figure 7), suggesting that the subclone has arisen to a very high frequency and is then obscured by the clonal mutations. Interestingly, 2/4 of these samples were MSI+, one plausible explanation is that the hypermutator phenotype results in an increased likelihood of acquiring an adaptive mutation when the tumour is still small enough for the clone to expand to a detectable frequency in a reasonable amount of time.

### Selection in constant size populations is more efficient

The analysis above considered only exponentially growing populations, which is a growth-pattern well supported by empirical data in many cancer types ^16−21^ Some tumours however, especially benign lesions, may reach a plateau in their growth, and consequently are better represented by sigmoidal-type growth models^22,23^. In a sigmoidal model of tumour growth, the tumour population at late times can be approximated as a population of constant size with continual turnover of cells. Interestingly, in a fixed size population it has been shown that the fixation time of beneficial mutations is proportional to the logarithm of the population size (see methods)^24^, which suggests that clonal expansions can be relatively rapid when the population is no longer growing.

To examine the effect of the population growth profile on subclone evolution, we simulated a model of fixed population size using a Moran process, and compared the speed with which subclones expand to the exponential growth model described above (figure 4A&B). The fitness advantage of a mutant in both fixed and growing populations was defined as the average offspring per generation (of the background host population). We introduced a fitter mutant in the growing population when the population was of size N, and simulated the Moran model for fixed size N; thus a new mutant starts out at a frequency 1/N in both cases. We followed the average frequency of the mutant over time. In the fixed population model the fitter mutant spreads through the population at a significantly faster rate (figure 4C; p<0.001), and we noted that subclonal expansions can also lead to subclonal clusters in the VAF distribution in a fixed population (figure 4E). We note that a constant population of cells that acquires new passenger mutations and undergoes neutral drift (figure 4A) results in a neutral tail in the VAF distribution that however, does not directly encode the mutation rate^6^ (figure 1B).

**Figure 4.**
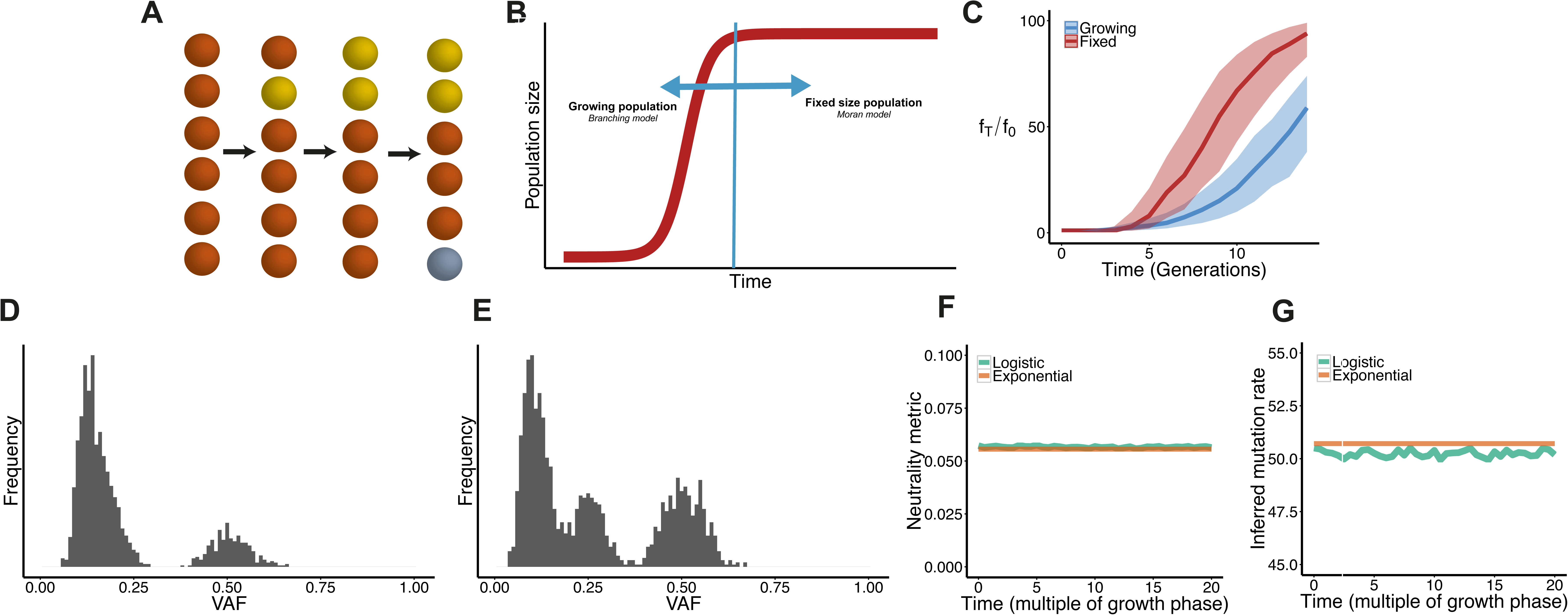
We used a Moran model to compare the dynamics between fixed size populations and growing population, **A**, and found that in fixed size population selection can be more rapid **C**. We simulated a moran model with N=100, and introduced a mutation at N=100 in the growing population so in both models the initial frequency of the mutation f_0_=1/100, clone has fitness advantage 1+s=0.5. Then measured the frequency at a later time f_T_, in the fixed population size the ratio f_T_/f_0_ increases quicker than in the growing population (p<0.001). The moran model can also produce VAF histograms similar to the neutral case, **D** (no selection, 300 generations) and the non neutral case **E** (1+s = 2, number of generations = 10). However simulating a tumour that grows logisitically and transitions into a moran model **B**, even when the population followed a moran model for 20 times longer than it was in the growth phase, the main signature of the VAF distribution is that of exponential growth given we observe no differences in our neutrality metric **F**, or the inferred mutation rate **G**.

Under a logistic regime, initial cancer growth is exponential, slowing to a constant population size (with turnover) once a ‘carrying capacity’ is reached. We investigated how this pattern of population growth influenced the measurement of evolutionary dynamics. We simulated logistic growth where the population first grows exponentially and then transitions into a Moran model (figure 4B). We found that assuming for example a small carrying capacity of 10^4^ cells, even if the fixed population size phase is 20 times longer than the growth phase, the dominant signature is that of the initial (neutral) growth, not the neutral drift within the fixed size population (figure 4F). Consequently, the mutation rate estimates match those measured in a purely exponential neutral model (figure 4F&G). We note that this is because tumours are very large populations, and effects of neutral drift during the constant phase are unlikely to be significant since the time it takes for variants to rise to a detectable frequency under these conditions is proportional to the population size *N*. Hence, even for barely-detectable tumours of 10^6^ cells, it would take approximately a million generations before seeing those drift tails: much longer than a human lifetime. Hence in cancer data, irrespective of whether or not the population has become constant, the VAF distribution encodes *initial* tumour growth, and neutral tails *do* accurately inform on the mutation rate.

Another growth regime that has been shown to be potentially applicable to cancer is power law growth. We note that such power law (boundary driven) growth leads to a slightly different scaling form for the VAF distribution (see supplementary note). We note that since only peripheral cells can proliferate in a power law model, the biological interpretation of neutral evolution in this growth regime is unclear.

## Discussion

We have demonstrated how the distribution of mutations in a tumour can be used to directly measure the evolutionary dynamics of subclones. We confirmed that subclonal selection causes an overrepresentation of mutations within the expanding clone, manifested as an additional ‘peak’ in the VAF distribution, in-line with the detection of subclonal clusters by many recent studies^7^. We note that our analysis predicts that even when subclonal selection plays a prominent role in tumour progression, the tumour will still show an abundance of low frequency variants (a 1/f-like tail, though the precise shape of the tail may be altered). This is a natural consequence of the tumour being a growing population, wherein the number of new mutations in the population is proportional to the population size.

Remarkably, our quantitative measurement of the size of the selective coefficient (relative fitness) of an expanding subclone in a tumour revealed that (large) subclones had experienced fitness advantages in excess of 20% greater than the ‘resident’ populations of the tumour. These fitness advantages are more than an order-of-magnitude greater than a previous estimate of 0.04%^25^. However, such large fitness increases are not unprecedented in somatic evolution: a study of the competitive advantage of mutant stem cells in the mouse intestine (a constant population size) showed that KRAS and APC mutant stem cells have a ~2-4 fold increase in the probability of fixing in the crypt^26^, and *TP53* mutant cells in mouse epidermis exhibited a 10% bias toward self renewal^27^. Moreover, our measured fitness increases are comparative with the most extreme values observed in experimental evolution settings, wherein most positively selected variants confer small percentage increases fitness^28,29^. Furthermore, a classical test for selection, the ratio of non-synonymous to synonymous variants (dN/dS) reveals small subset of genes (<20 in a pan-cancer analysis) with extreme dN/dS values indicative of strong selection^30^.

Importantly, our analysis shows that there can be heterogeneity in the evolutionary process within a tumour: four regions of a single lung cancer were found to be evolving neutrally whereas an additional region showed strong evidence of subclonal selection. We note that our analysis does not *ipso facto* identify the cause of the subclonal expansions. Irrespective, we note that any change experienced by a subclone that results in increased fitness, including copy number variation, epigenetic changes, point mutations or cell-extrinsic effects (clonal interactions or microenvironment effects) will be ‘read out’ as causing selection in the VAF distribution. This is because selection is inferred using only the frequency of SNVs, which will shift in frequency due to hitchhiking, regardless of the underlying mechanism.

We note that our analysis indicates that even if cancer subclones experience pervasive weak selection, that this weak selection does not cause the VAF distribution to deviate detectably from the distribution expected under strict neutrality. Thus, our analysis implies that so-called ‘mini-drivers’ could well be common in cancer^31^, but that each mini-driver has a corresponding ‘mini’ effect on the subclonal composition of a tumour, and correspondingly that dramatic changes to the tumour population are only caused by ‘major-drivers’. Additionally, we note that for a cancer that is experiencing prevalent weak selection, neutrality provides an entirely adequate description of the evolutionary dynamics as measured by moderate depth sequencing data.

The noise inherent in the data means that measuring evolution in the cancer genome is extremely challenging. For this reason we concentrated our efforts on a small number of deeply-sequenced tumours, as the depth of sequencing in particular has a large effect on the ability to resolve subclonal structure in the genome (supplementary figure 8). We acknowledge that features that are not described in our model, principally the spatial structure of the tumour, could effect the accuracy of our estimates of evolutionary parameters^32^. Spatial models of tumour evolution can help elucidate other important biological parameters such as the degree of mixing within tumour cell populations^10^, which is a purely spatial phenomenon and cannot be quantified using non spatial models such as ours. Multiple samples per tumour also increase the power to detect selection within a cancer, as the probability that a ‘clone boundary’ where selection is evident will be sampled is increased.

In summary, we have shown how clonal selection shapes the frequency distribution of subclonal mutations within a tumour, and used this knowledge within a mathematical framework to directly measure, *in vivo* in human malignancies, the fundamental evolutionary parameters that control subclonal evolution. These data give new insight into the process of human carcinogenesis, and show the power of a quantitative phenomenological framework for understanding cancer evolution.

## Methods

### Simulating tumour growth

We implemented a branching process simulation of cell divisions during tumour growth, followed by a sampling scheme that recapitulates the characteristics of cancer sequencing data. Cancer sequencing data is plagued by various sources of noise, so this final step is required to ensure that the underlying evolutionary dynamics that govern cancer growth are not confounded by the noisy signal. First we will introduce the simulation framework for an exponentially expanding population where all cells have equal fitness. Later we show how elements of the simulation can be modified to include differential fitness effects of cells and non-exponential growth.

Tumour growth begins with a single transformed cancer cell that has acquired the full set of genetic alterations necessary for malignancy. This first cell will therefore be carrying a set of mutations (the number of these mutations can be modified), which will be present in all subsequent lineages and thus are clonal (present in all cells) in the population. Any subsequent mutations that are acquired are likely to remain subclonal, but due to the stochastic nature of the model there are scenarios - such as a slow growing population with high cell death - where the mutations that are acquired during the first few divisions can become clonal. Expressions for the probability that a subclonal mutation becomes clonal have been derived elsewhere^33^.

The dynamics of tumour growth are governed by a birth rate and death rate that are set at the beginning of the simulation. These can be modified to include selection and non-exponential growth. Given a birth rate *b* and death rate *d* (*b*>*d*, for a growing population), the average population size at time t will be given by,

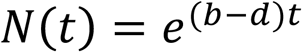

We set *b*=*log(2)* for all the simulations, such that in the absence of cell death the population will double in size at every unit of time. The tumour grows until it has reached a specified size *N_final_*, where the simulation stops. At each division, cells acquire υ new mutations, where υ is drawn from a Poisson distribution with mean *μ*. We assume new mutations are unique (infinite sites approximation). Not all divisions will result in surviving lineages, the probability of a cell division producing a surviving lineage, *β* can be written as the following in terms of the birth and death rates

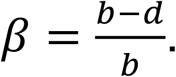

### Selection

To include the effects of selection, a mutant is introduced into the population that grows at a faster rate than the host population. We only consider the cases of one or two subclonal population under selection. The number of large-effect driver mutations in a typical cancer is thought to be small (<10 see ref. ^34^), so this restriction was made for pragmatic reasons. Fitter mutants can have a higher birth rate, a lower death rate or a combination of the two, all of which results in the mutant growing at a faster rate than the host population. Given that the host/background population has growth rate *b_H_* and death rate *d_H_*, and the fitter population has growth rate *b_F_* and death rate *d_F_* we define the selective advantage s of the fitter population as:

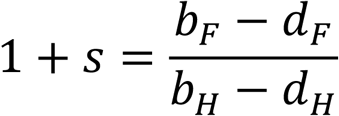

Fitter mutants can be introduced into the population with a specified advantage s and at a chosen time *t_1_*, allowing us to explore the relationship between the strength of selection and the time the mutant enters the population.

A number of simplifications to our simulation scheme were made to improve computationally efficiency. This is particularly relevant for the Bayesian inference approach that requires many millions of individual simulations to be performed.

The first simplification neglected cell death, and so models differential subclone fitness by varying the birth rate only. Setting the death rate to 0 (e.g *β* = 1, all lineages survive) increases simulation speed because a smaller number of time steps are required to reach the same population size.

This simplification affects our ability to measure the effective mutation rate, 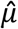, which is the true mutation rate *μ*, divided by the probability of having 2 surviving offspring

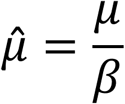

The effective mutation rate is encoded in the low-frequency (1/f-like) tail of the distribution. In the presence of one or more subclones, the low-frequency tail consists of a combination of two or more 1/f tails. If there are large differences in the *β* value between subclones, then the inference on the effective mutation rate from the gradient of the low-frequency tail may be incorrect. However this is true only if there are large differences in *β* in the different subclones. To show this, we simulated subclones with a range of different *β* values, and inferred the mutation rate from the low frequency tail. Even in cases where the death rate was very different in the subclone compared to the host population (*β* = 1.0 vs *β* = 0.5) the mean error on the estimates of the mutation rate was 42% (supplementary figure 9) - e.g. significantly less than the order of magnitude previously measured between cancer types ^6^.

The second simplification restricts simulations to only a small population size. We note that the VAF distributions hold no information on the population size, meaning that a simulation can produce VAF distributions that match real data even when the simulated population size is unrealistically small. Moreover, subclone fitness and size are both measured relative to the (unmeasurable) overall fitness and size of the entire tumour population. For example, imagine a tumour growing at some (exponential) rate, which could be either slow or fast. Within the growing tumour, a subclone forms. To achieve its final size, in the fast growing tumour the subclone must grow significantly faster than the host population for a short time, or relatively slowly for a long time. In comparison, within the slower growing tumour, the subclone could grow comparatively slower for a shorter period and still achieve the same final size. In other words, both the time elapsed since the formation of a subclone and the final size of the subclone in the tumour together scale with the (exponential) growth rate and final size of the tumour as a whole (with the scaling specified in equations [3] & [5]). Therefore we can simulate small tumours wherein subclones have large fitness advantages, and then scale our estimates of the selective advantage using realistic population sizes and growth rates to obtain biologically meaningful estimates of the evolutionary parameters. The size of the simulated tumour has no impact on the accuracy of parameter inference, as long as we simulate for a time long enough for any possible subclones to accumulate enough mutations to be consistent with the data.

To appropriately scale the estimates of s requires inputting an estimate of the age of the tumour in terms of tumour doublings into equation [5]. Assuming a final population size of *N_end_*, we can calculate *t_end_* as,

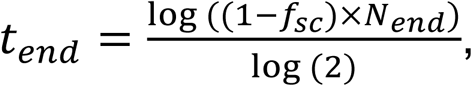

where *f_sc_* is the frequency of the subclone. We assumed *N_end_* = 10^9^, for generating the posterior distributions in figure 3. We also generated posterior distributions for s as a function of *N_end_*, for all samples that showed evidence of subclones, supplementary figure 10.

To demonstrate the validity of this approach, we simulated a comparatively large tumour (10^5^) with a high death rate (*β* = 0.25) and a subclone with a lower death rate (*β* = 0.74), and then used our inference scheme (see below) with beta=1 for both residual and subclonal cells (e.g. no death) to attempt to recover the selective advantage of the subclone in our simulated tumour. The posterior distributions were correctly centred around the true parameters(Fig 2).

### Simulation method

A rejection kinetic Monte Carlo algorithm was used to simulate the model ^35^ Due to the small number of possible reactions (we consider at most 3 populations with different birth and death rates) this is more computationally efficient than a rejection-free kinetic Monte Carlo algorithm such as the Gillespie algorithm. The input parameters of the simulation are given in table 1

**Table 1:** Input parameters for simulation

|  |  |
| --- | --- |
| **b** | Birth rate of host population |
| **d** | Death rate of host population |
| **b<sub>F</sub>** | Birth rate of fitter populations, each new population will have a unique b<sub>F</sub> |
| **d<sub>F</sub>** | Death rate of fitter populations, each new population will have a unique d<sub>F</sub> |
| **s** | Selective advantage of fitter populations calculated from b<sub>F</sub> and d<sub>F</sub> |
| **μ** | Mutation rate |
| **t<sub>event</sub>** | Time when fitter mutant is introduced |
| **N<sub>final</sub>** | Maximum population size, simulation stops once this is reached |

The simulation algorithm is as follows:

1. Simulation initialized with 1 cell and set all simulation parameters
2. Choose a random cell, i from the population
3. Draw a random number r~Uniform(0, b_max_+d_max_), where b_max_ and d_max_ are the maximum birth and death rates of all cells in the population.
4. Using r, cell i will divide with probability proportional to its birth rate b_i_ and die with probability proportional to its death rate d_i_. If b_i_+d_i_ <b_max_+d_max_ there is a probability that cell i will neither divide nor die. If *β* = 1, ie no cell death then in the above d_max_ = 0.
5. If cell divides, daughter cells acquire *v* new mutations where *v* ~Poisson(μ)
6. Time is increased by a small increment 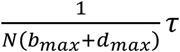, where *τ* is an exponentially distributed random variable ^36^
7. Go to step 2 and repeat until population size is N_max_

The output of the simulation is a list of mutations for each cell in the final population.

### Sampling

To mimic the process of data generation by high-throughput sequencing we performed various rounds of empirically-motivated sampling of the simulation data. Sequencing data suffers from multiple sources of noise, most importantly for this study is that mutation counts (VAFs) are sampled from the true underlying frequencies in the tumour population (both because of the initial limited physical sampling of cells from the tumour for DNA extraction, and then due to the limited read depth of the sequencing). Additionally it is challenging to disentangle mutations that are at low frequencies from sequencing errors and consequently only mutations above a frequency of around 5-10% for 100X sequencing are detectable ^11^. The ability to resolve subclonal structures is dependent on the depth of sequencing. This is shown in supplementary figure 8, where the same simulation has been sampled to different depths and the subclonal architecture is progressively obscured as the depth decreases.

For mutation *i* the frequency of mutation is binomially distributed *f_t_*~*B*(*n* = *D_i_*,*p* = *VAF_true_*), where the sequencing depth D is itself a binomially distributed random variable and *VAF_true_* is the known VAF of the mutation before sampling. The “sequenced” VAF is thus 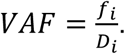 Sequencing data is often found to be overdispersed, for cases where we found the data to be overdispersed we used the Beta-Binomial distribution ^37,38^. In this model the frequency of mutation i is distributed according to *ƒ_i_*~*BetaBin*(*n* = *D_i_*,*p* = *VAF_true_*,*ρ*) where *ρ* is the degree of overdispersion and introduces additional variance to the sampling. For *ρ* = 0, the model is the usual Binomial model. All subsequent analysis is then done using these resultant sequencing noise processed VAF distributions.

### Testing Neutrality

To assess what evolutionary parameters of selection lead to an observable deviation from neutrality we devised multiple metrics to detect deviations from the prediction of the neutral model. Previously we showed that under neutrality, the distribution of mutations with a frequency greater than f is given by ^6^:

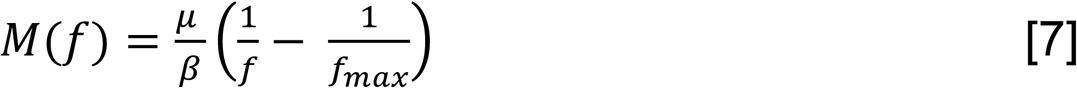

Previously, we fit a linear model of M(f) against 1/f and used the R^2^ measure of the explained variance as our measure of the goodness of fit.

Another approach is to use the shape of the curve described by equation [7] and test whether our empirical data collapses onto this curve. To implement this, here we introduce a *universal neutrality curve*, *M̅*(*ƒ*). Given an appropriate normalization of the data, any mutant allele frequency distribution governed by neutral growth will collapse onto this curve. We can normalize the distribution described by equation [7] by considering the maximum value of M(f), which is given when f=f_min._

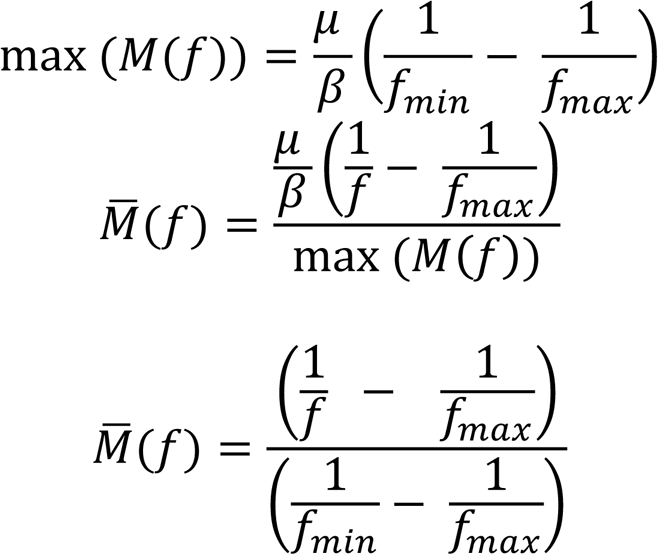

*M̅*(*f*) is independent of the mutation rate and the death rate, which allows comparison with any dataset. To compare this theoretical distribution against empirical data we used the Kolmogorov distance, D_k_, the Euclidean distance between *M̅*(*f*) and the empirical data and the area between *M̅*(*ƒ*) and the empirical data. The Kolmogorov distance D_k_ is the maximum distance between two cumulative distribution functions. Mathematically D_k_ for *M̅*(*f*) and an empirical cumulative distribution - *Ĝ*(*f*) is defined as

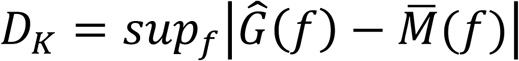

where sup is the supremum of the set of distances. Supplementary figure 11 provides a summary of the different metrics.

To assess the performance of the 4 classifiers we ran 10^5^ neutral and nonneutral simulations and compared the distribution of the metrics for these two cases. Due to the stochastic nature of the model, not all simulations that include selection will result in subpopulations at a high enough frequency to be detected, therefore to accurately assess the performance of our tests we only included simulations where the fitter subpopulation was within a certain range (20% and 70% of the final tumour size). All 4 metrics showed significantly different distributions between neutral and non-neutral cases (supplementary figure 1). Under the null hypothesis of neutrality and a false positive rate of 5%, the area between the curves was the metric with the highest power (67%) to detect selection, slightly outperforming the Kolmogorov distance and euclidean distance, with the R^2^ metric showing the poorest performace with a power of 61% (table S1 and supplementary figure 1).

We also plotted receiver operating characteristic (ROC) curves by varying the discrimination threshold of each of the neutrality tests and calculating true positive and false postive rates (using a dataset derived from simulations with subclonal populations at a range of frequencies, supplementary figure 2). This also showed that the R^2^ had the least disriminatory power, with the other 3 performing equally well (see table S2 for AUC). Increasing the range of allowed subclone sizes decreased the classifier performance, likely because the subclone could merge into the clonal cluster or 1/f tail when it took a more extreme size.

### Statistical Inference

We used Approximate Bayesian Computation (ABC)^39^ to infer the evolutionary parameters in our stochastic tumour evolution model that produced variant allele frequency distributions consistent with real sequencing data. We also validated the accuracy of our inferences using simulated sequencing data where the true underlying evolutionary dynamics was known.

As in all Bayesian approaches, the goal of the ABC approach was to produce posterior distributions of parameters that give the degree of confidence that particular parameter values is true, given the data. Given parameter vector of interest θ and data D, the aim was to compute the posterior distribution 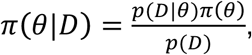 where *π*(*θ*) is the prior distribution on θ and *p*(*D*|*θ*) is the likelihood of the data given θ. In cases where calculating the likelihood is intractable, as was the case here where our model cannot be expressed in terms of well known and characterized probability distributions, approximate approaches must be sought. The basic idea of these ‘likelihood free’ ABC methods is to compare simulated data, for a given set of parameter values, with observed data using a distance measure. Through multiple comparisons of different input parameter values, we can produce a posterior distribution of parameter values that minimise the distance measure, and in so doing accurately approximate the true posterior. The simplest approach is called the ABC rejection method and the algorithm is as follows^40^:

1. Sample candidate parameters θ^*^ from prior distribution π(θ)
2. Simulate tumour growth with parameters θ^*^
3. Evaluate distance, δ between simulated data and target data
4. If δ ≤ ε accept parameters θ^*^
5. If δ > ε accept parameters θ^*^
6. Return to 1

We used an extension of the simple ABC rejection algorithm, called Approximate Bayesian Computation Sequential Monte-Carlo (ABC SMC) ^12,41^. This method achieves higher acceptance rates of candidate simulations and thus makes the algorithm more computationally efficient than the simple rejection ABC. It achieves this by propagating a set of ‘particles’ (sample parameter values) through a set of intermediate distributions with ever decreasing ε until the target ε_T_ is reached, using an approach known as sequential importance sampling^42^. The ABC SMC algorithm also allows for Bayesian model selection to be performed by placing a prior over models and performing inference on the joint space of models and model parameters, (m, θ_m_). In contrast to many applications of ABC that use summary statistics, we use the full data distribution, thus avoiding issues of inconsistent Bayes factors due to loss of information^43,44^. For further details on the algorithm see references^41,45^ and the supplementary note on the specific details of our implementation. Bayes factors for all data are shown in table S3.

We used a modified version of the Kolmogorov distance as our distance function which has been used in similar inference problems^46^. The well-known Kolmogorov distance is however invariant to the mutation rate, one of the parameters we would like to infer. We therefore use an unnormalized version of this statistic that will depend on the mutation rate. Simply, we calculate M(f) for both datasets and as in the Kolmogorov distance take the maximum distance between the experimental data and the synthetic data.

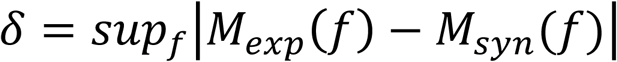

We only perform the fit for mutations with VAF>f_min_, where f_min_ is the detection threshold of the data which we deem to be the point at which the 1/f peak tails off at low frequency. The priors used for inference are shown in the supplementary note.

### Bioinformatics analysis

Where available we used variant calls provided from the original studies. The processing of the lung cancer sequencing data^14^ and gastric cancer^6,15^ is explained elsewhere. We additionally applied the Sequenza algorithm^47^ to infer allele specific copy number states and estimate the cellularity. Copy number aberrations could also potentially result in the multi-peaked distribution we observe^48^, hence we only used mutations that were found in regions identified as diploid (and without copy-neutral LOH). The Sequenza algorithm also estimates the cellularity of the sample, which we used to correct the VAFs. For a cellularity estimate κ, the corrected depth for variant *i* will be *d*̅_i_ =*κ*×*d_i_*. Due to the computational cost of fitting our model with high mutation rates we randomly sampled 2000 mutations from the gastric cancer samples and performed the analysis with these mutations.

As noted our simulation can account for the over-dispersion of allele read counts. To measure the over-dispersion parameter *ρ*, we fitted a Beta-Binomial model to the clonal cluster where we know *VAF_true_* = 0.5. We used Markov Chain Monte Carlo (MCMC) to fit the following model to the right hand side of the clonal cluster so as to minimize the effects of the 1/f distribution or subclonal clusters:

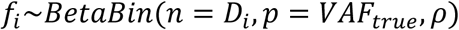

where *D_i_* is the sequencing depth, *f_i_* is the allele read count and *ρ* is the overdispersion parameter. We then used this estimate for *ρ* in the simulation sampling scheme. Supplementary figures 7 and 12 shows the fits to the clonal cluster for all our data using both the Beta-Binomial and Binomial model.

### Logistic Growth

In the logistic growth model, growth is density dependent and the environment has a maximum number of individuals it can support, which is commonly referred to as the carrying capacity, K of the population. The differential equation for logistic population growth is

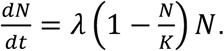

In the logistic population growth model, the birth and death rates of individuals in the population are proportional to the population size

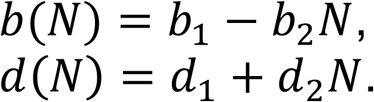

Where b_1_ and d_1_ are the intrinsic birth and death rates, and b_2_ and d_2_ can be calculated given a carrying capacity K from:

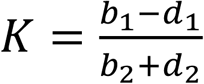

When b_2_ = d_2_ = 0, we recover exponential growth.

We consider two models of logistic growth, one where the birth rate decreases as the population grows (d_2_=0), and the other when the death rate increases (b_2_=0) as the population grows. This lets us explore the importance of stochastic effects. In the second model when b_2_=0, there is a fast turnover in cells, while in the first model turnover is slow. We used b_2_=0 model for the simulations in figure 4.

### Moran Model

The Moran model is a classic model from population genetics, it is a stochastic birth death process where at each time step one individual is chosen to die and one is chosen to replicate ^49^ Individuals that have fitness advantages are more likely to be chosen to replicate, the selection coefficient is often defined as relative increase in the average number of offspring per generation: a fitter individual will on average have 1+s more offspring. It has been shown that the average fixation time (in generations) of a neutral mutation is ~ N. In the case of a beneficial mutation the time to fixation, T_fix_ is given by^50^

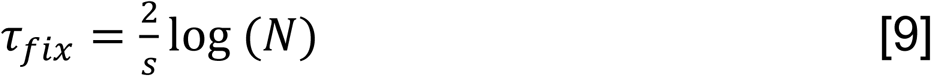

Therefore for a fixed size neutral population, the timescales over which mutations may rise to observable frequencies is likely longer than the age of the tumour, see table 2. Results consistent with our simulations that demonstrated that if a tumour follows a logistic growth model, the dominant signal in the VAF distribution is that of the early exponential growth (Fig 4). Selection however can results in mutations reaching observable frequencies rapidly.

**Table 2.** Fixation times in a neutral Moran model and a Moran model with selection.

| Model | Equation for $\tau_{fix}$ | s | N | $\tau_{fix}$ (generations) |
| --- | --- | --- | --- | --- |
| Selection | $\tau_{fix} \sim \frac{2}{s} \log(N)$ | 0.5 | $10^9$ | 82 |
| Neutral | $\tau_{fix} \sim N$ | 0.0 | $10^9$ | $10^9$ |

## Acknowledgements

We thank Weini Huang for fruitful discussions on selection in fixed size populations.

## Supplementary Material

**Figure S1.**
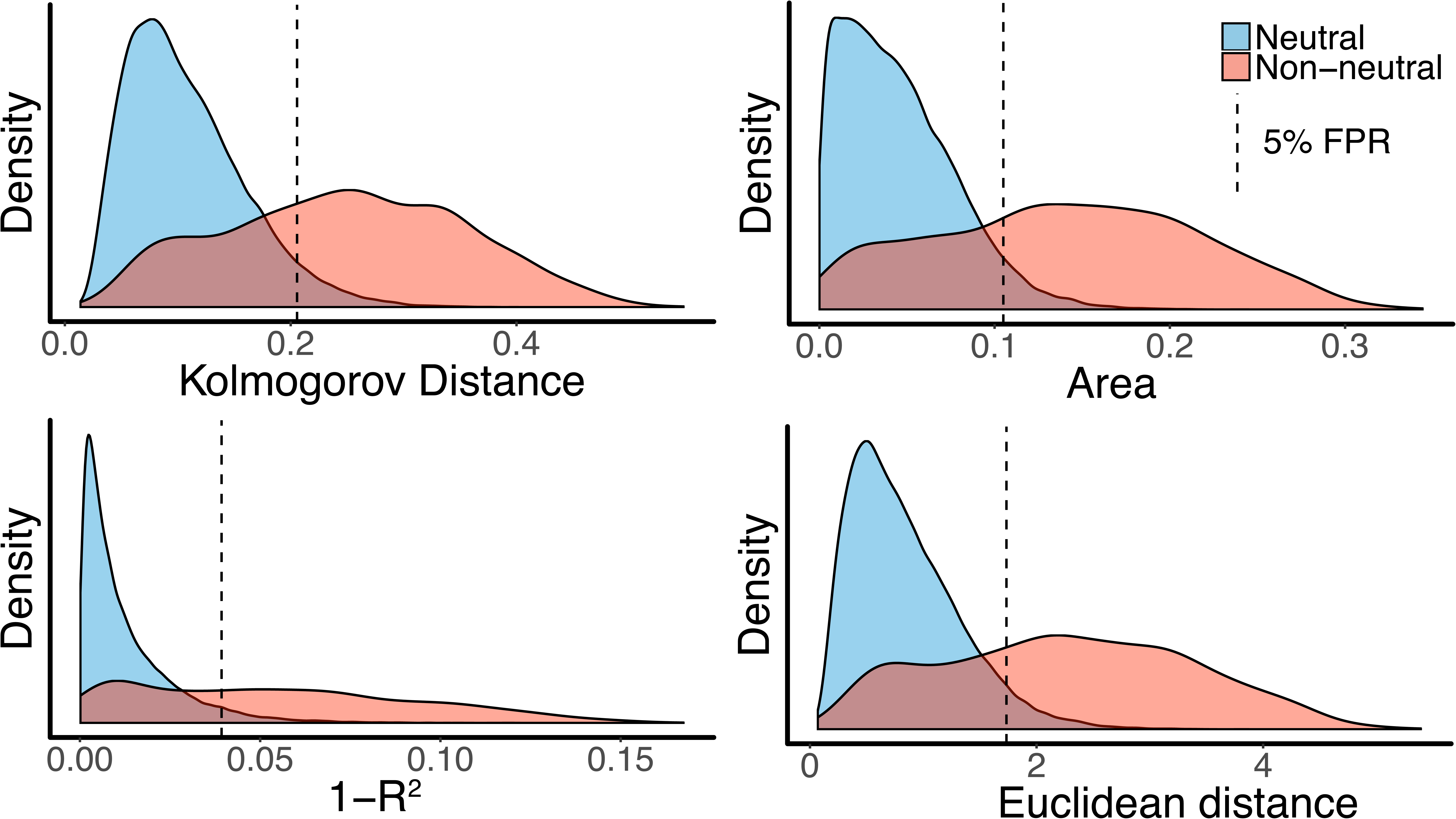
Taking 10^5^ neutral simulations and 10^5^ non neutral simulations (100X ‘sequencing’ depth) with a subclone with frequency greater than 20% and smaller than 70% we found that all metrics had significantly different distributions.

**Figure S2.**
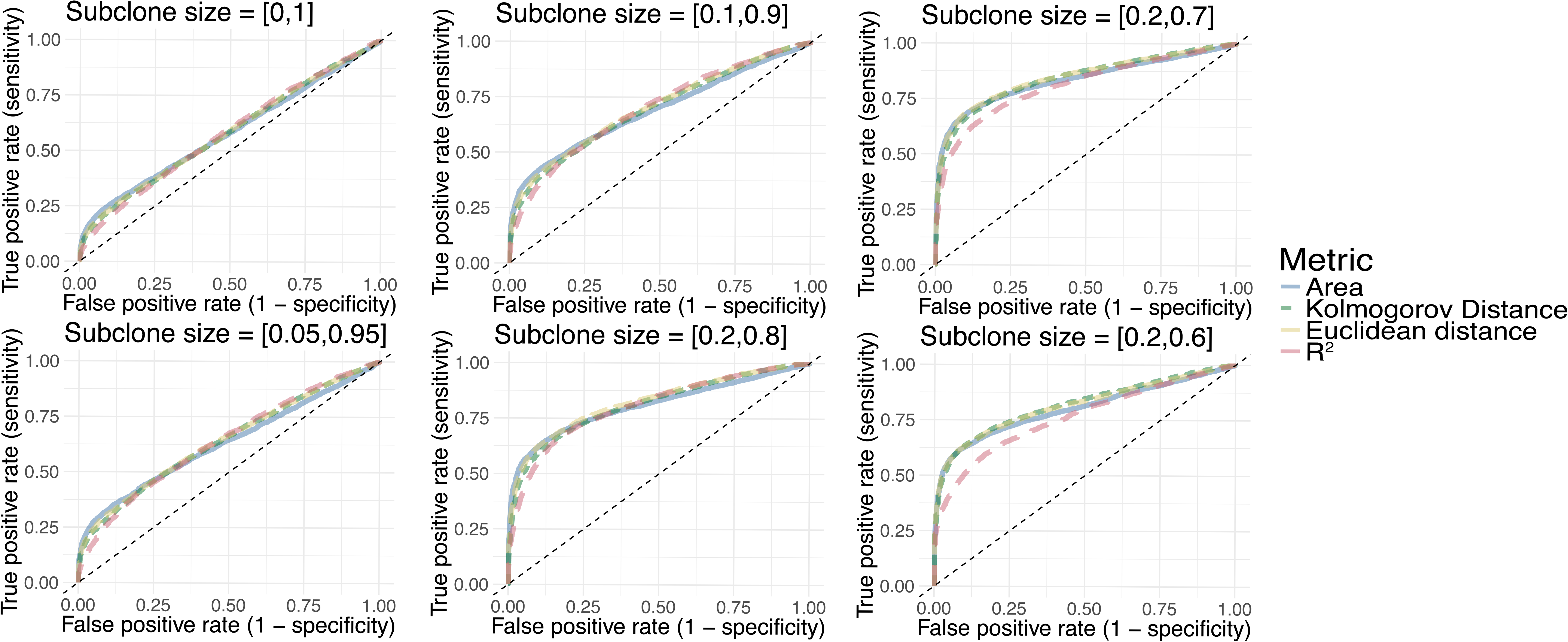
ROC analysis showed that the ability to detect deviations from the neutral model depends on the frequency of the subclone and that the area metrics is the most performant as it showed the largest area under the curve (see table S2 for values).

**Figure S3.**
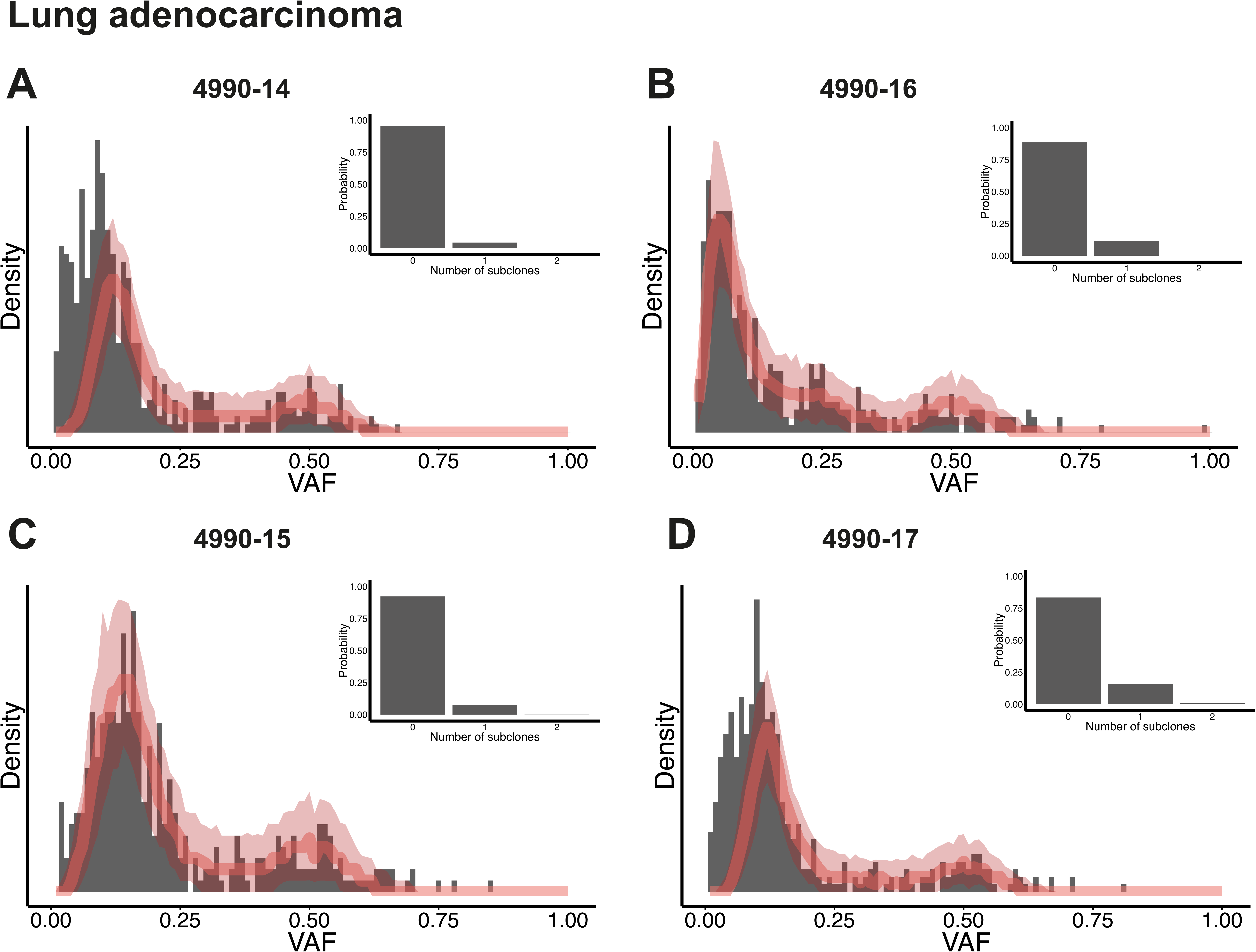
Applying our method to the 4 other samples from patient 4990 we found them to all be consistent with a neutral model with bayes factors in favour of the neutral model over the 1 subclone model ranging from 5.2 to 21.8 (see table S3).

**Figure S4.**
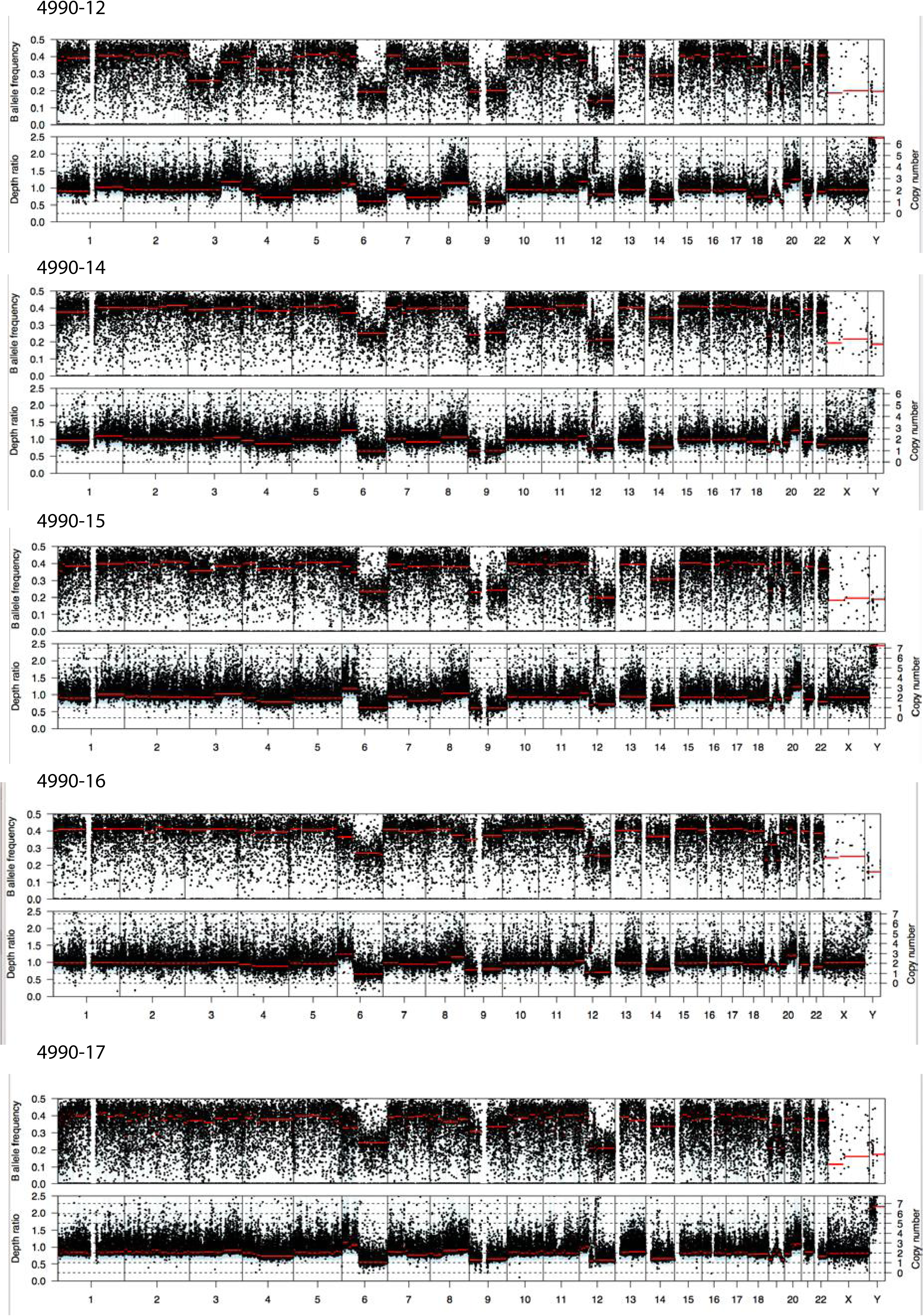
Copy number profiles for the 5 lung adenocarcinoma samples. Sample 4990-12 appears to have a CNA not present in the other samples.

**Figure S5.**
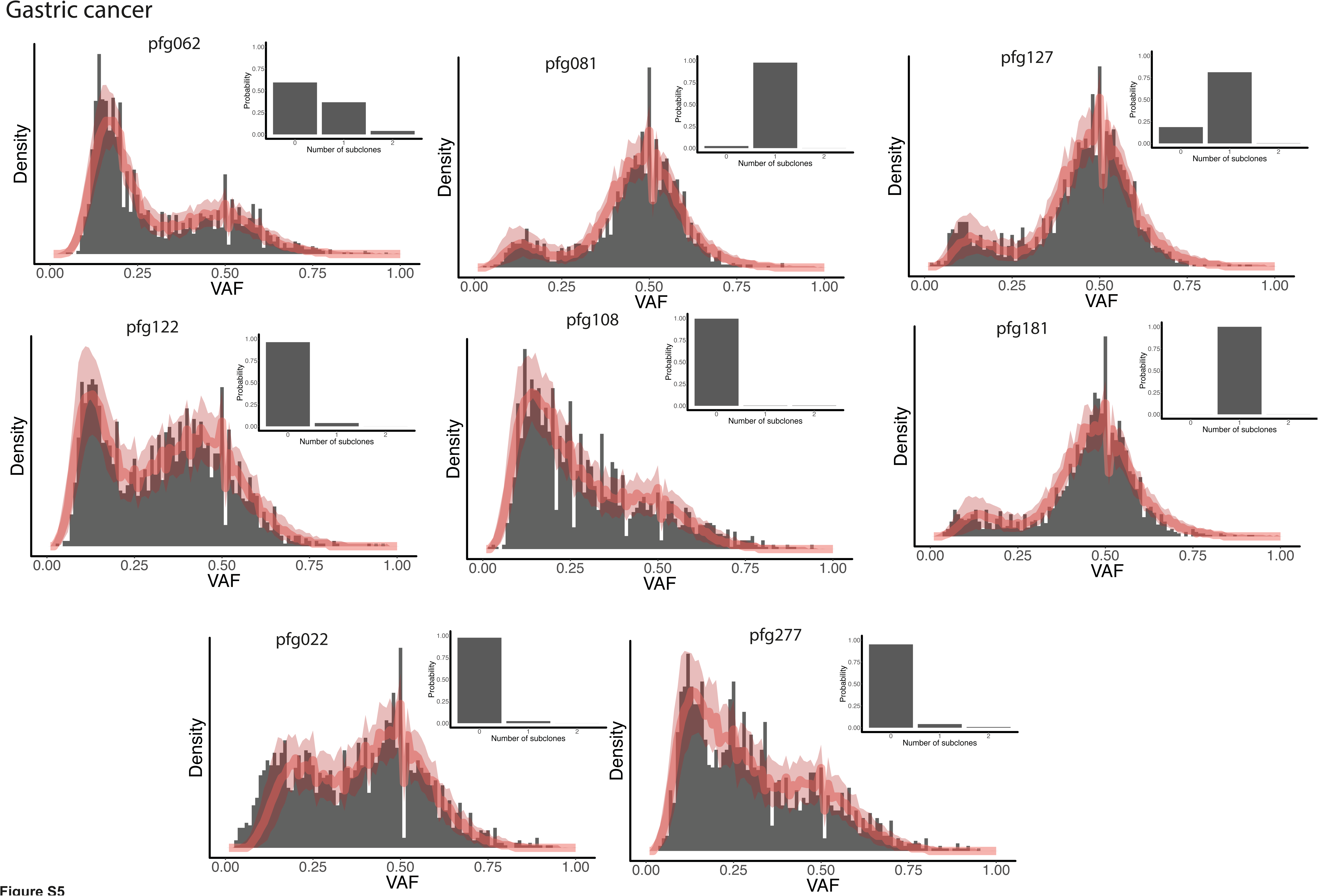
Inferred subclonal structure from 8 gastric cancers. 3 showed strong evidence of a subclonal population, while 5 were consistent with a neutral evolutionary model.

**Figure S6.**
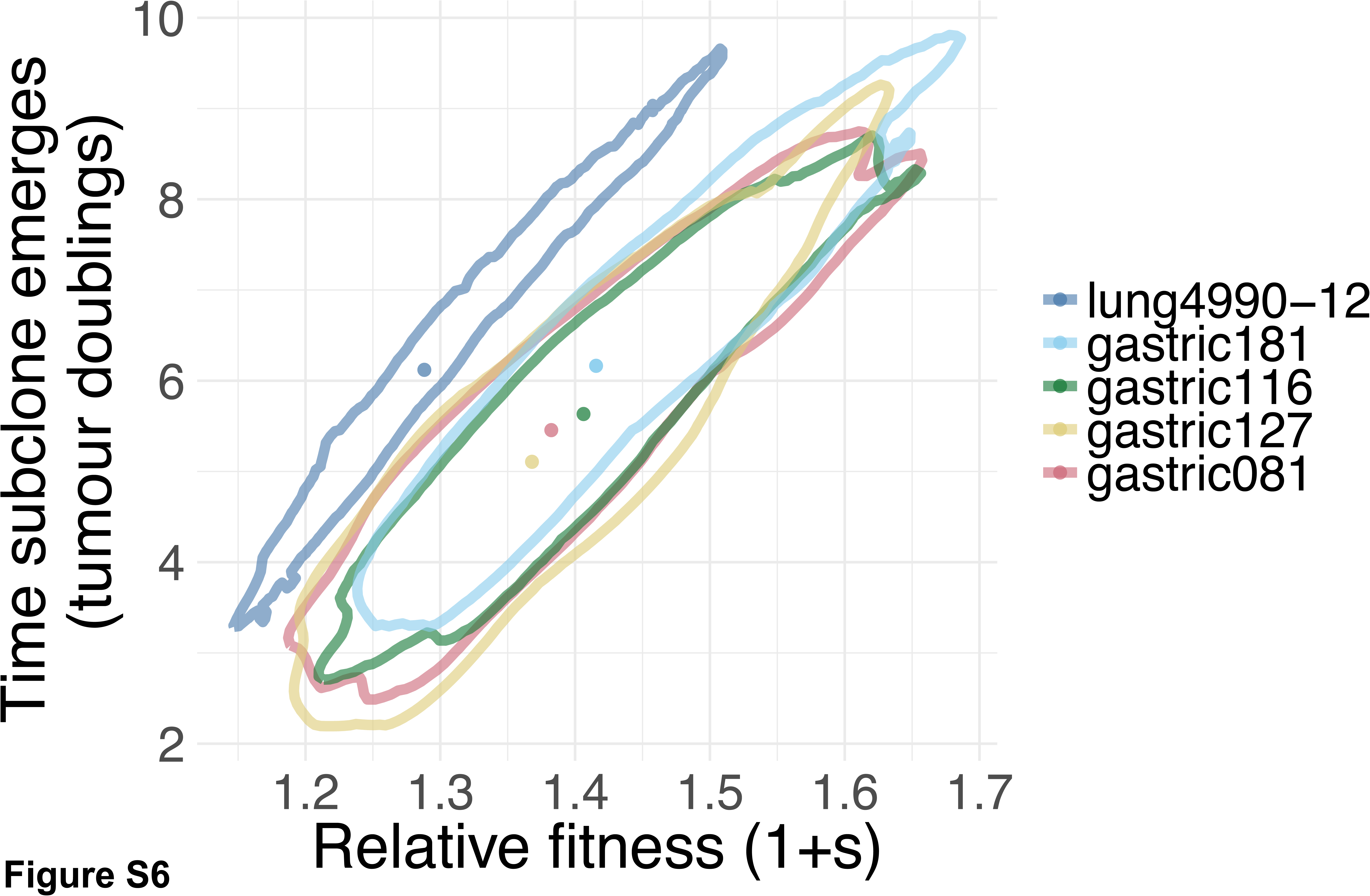
Time subclone emerged and selective advantage of subclones for the 4 samples where we identified subclonal population under differential selection. Points are median values and lines are 95% credible intervals.

**Figure S7.**
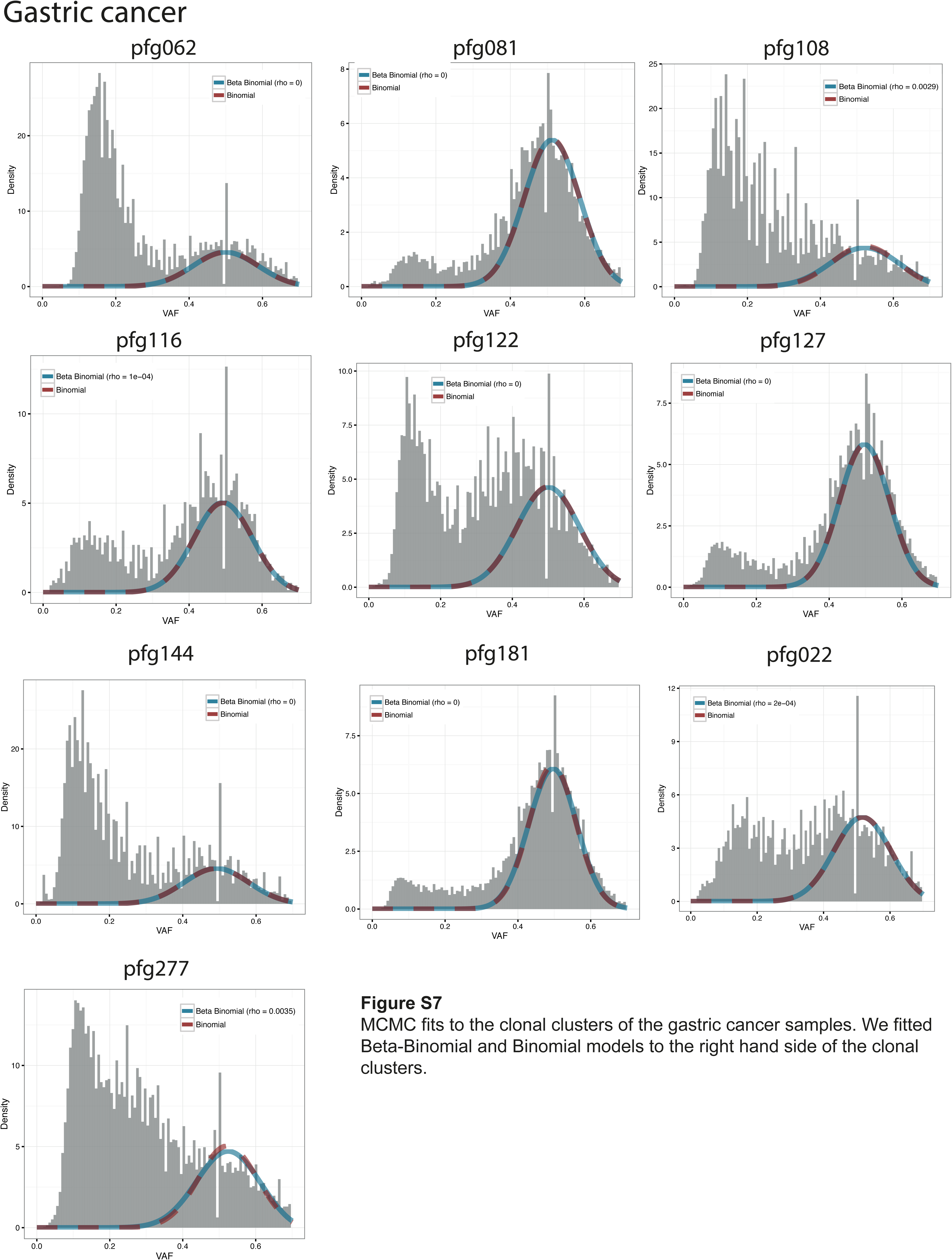
MCMC fits to the clonal clusters of the gastric cancer samples. We fitted Beta-Binomial and Binomial models to the right hand side of the clonal clusters.

**Figure S8.**
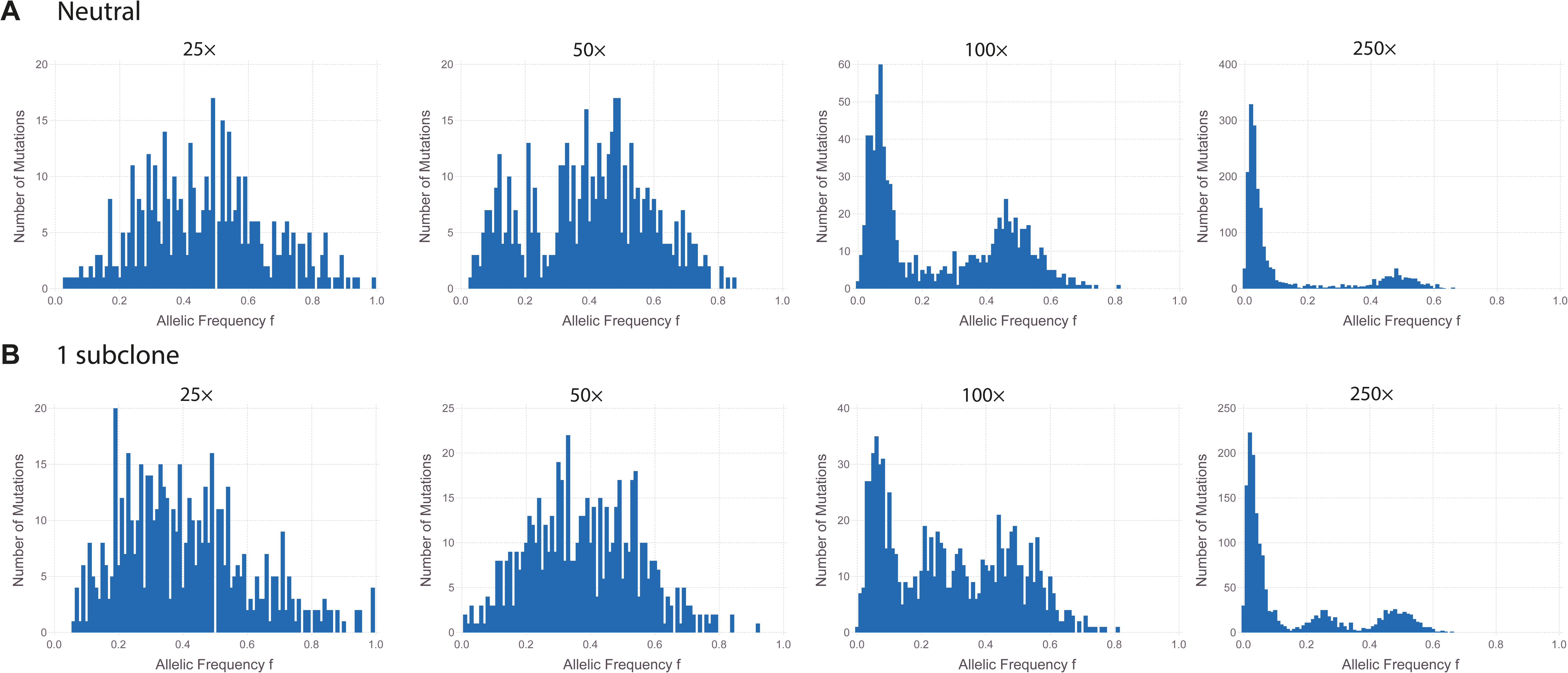
Simulated tumour under neutral growth in silico sequenced to 25X, 50X, 100X and 250X, **A**. Simulated tumour under non-neutral growth in silico sequenced to 25X, 50X, 100X and 250X **B**. Subclonal structure becomes more obscured as the depth of sequencing decreases. We required 5 “reads” to be observed for the variant to be detected. So the detection limit is 5/depth, so for 100X sequencing the limit is 5%.

**Figure S9.**
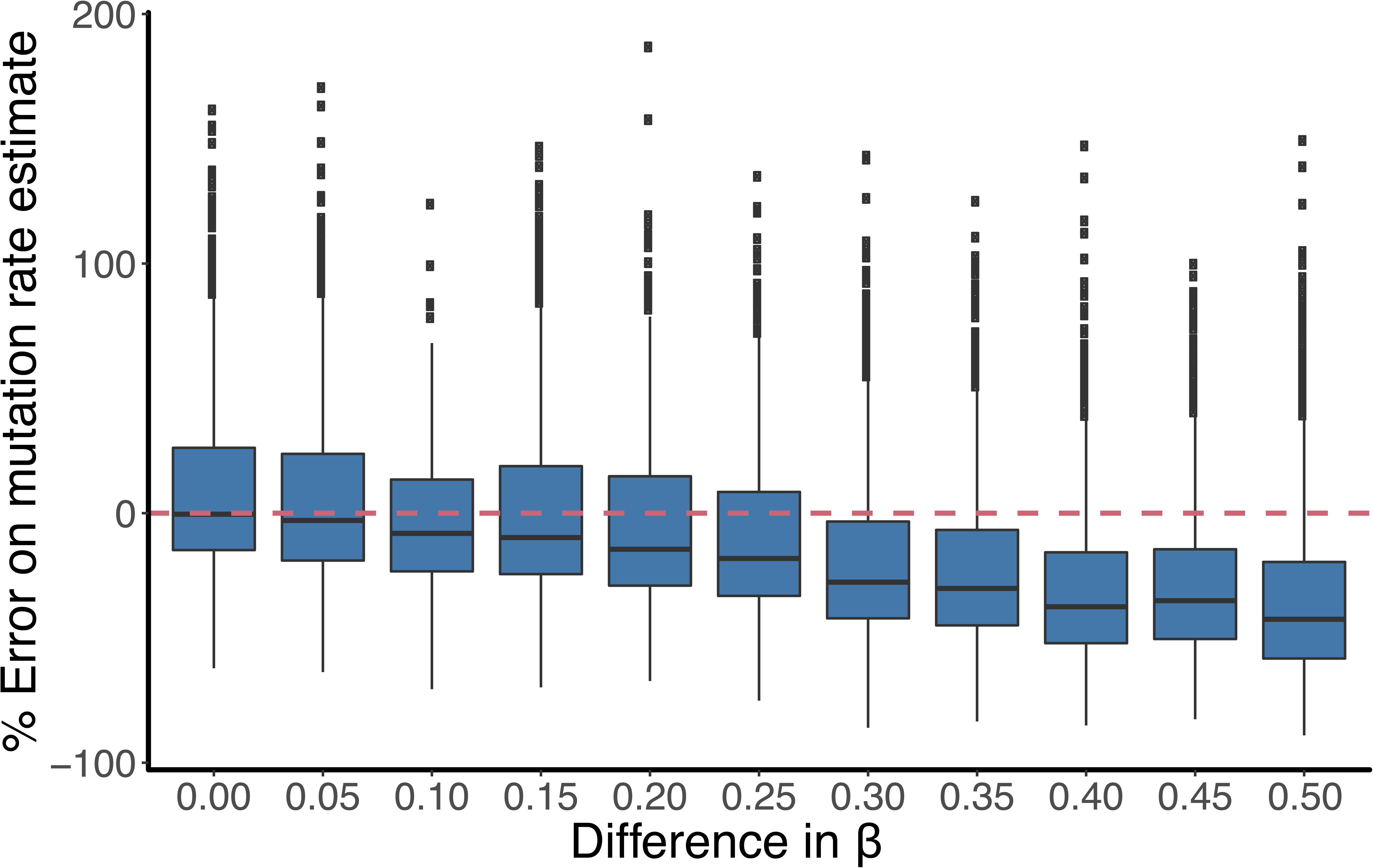
We ran a large number of simulations with a single subclone where the probability that a new lineage survives, β is different between the background host population and the subclone. We then measured the mutation rate by fitting a linear model to the left hand peak. The % error on the inferred mutation rate increases as the difference between β values increases but the mean error is not more than 50% even when Δβ=0.5.

**Figure S10.**
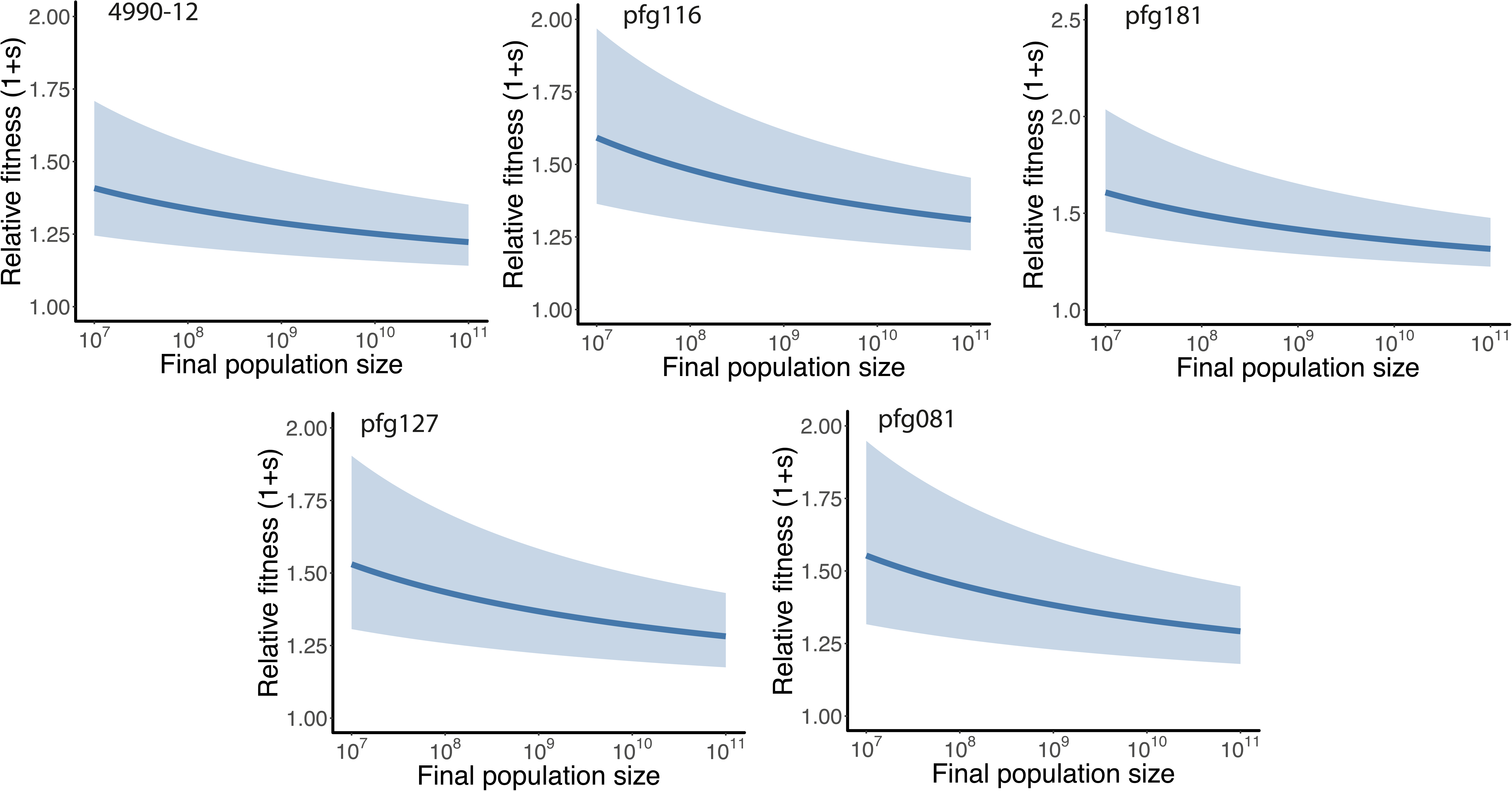
For all samples identified with a subclonal population, posterior distribution for the relative fitness as a function of the assumed final population size.

**Figure S11.**
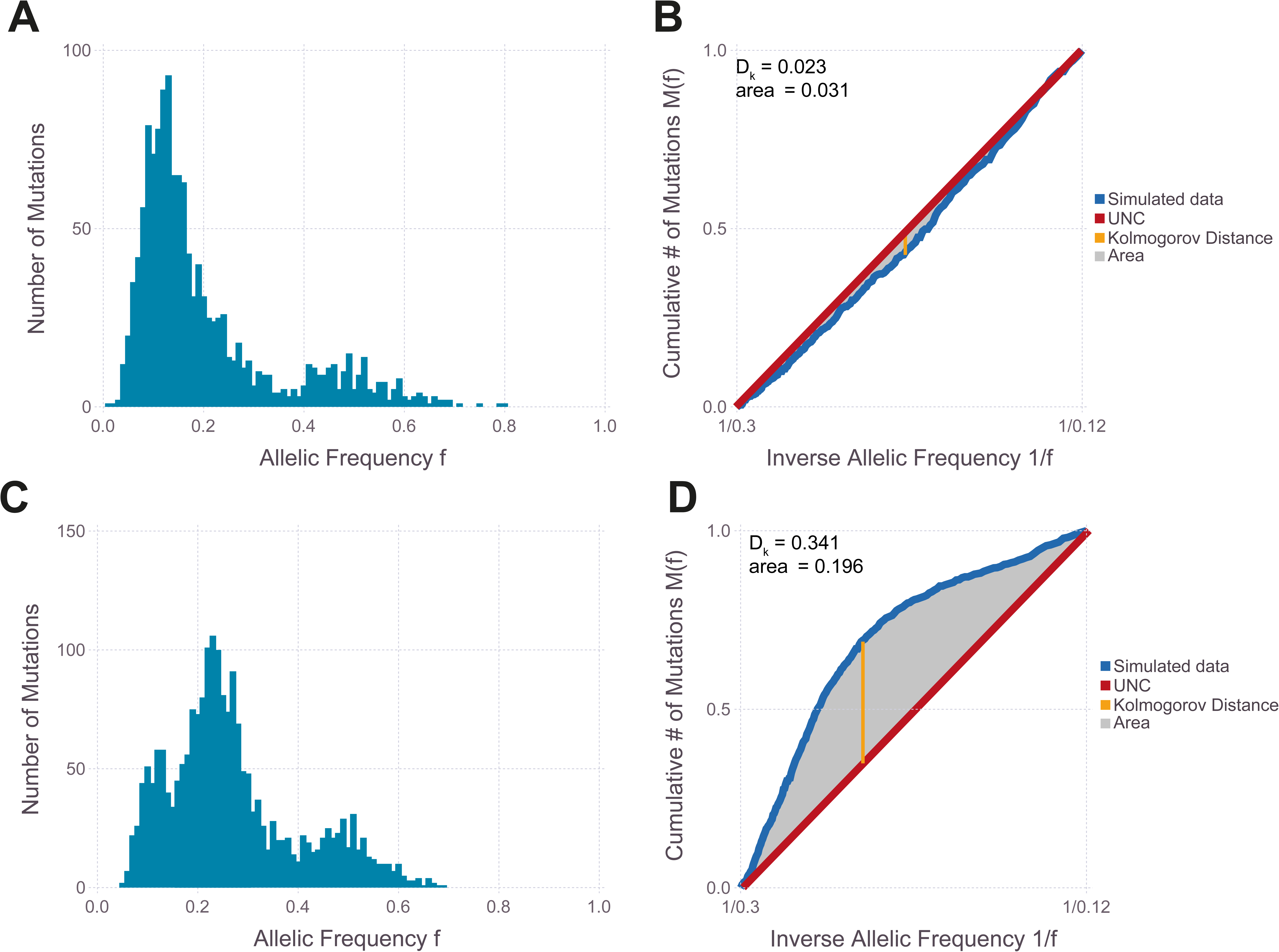
Simulated neutral simulation **A** and simulated simulation with 1 subclone **C**. To accept or reject the neutral model we tested a number of metrics where we compared the data (blue line) to the universal neutrality curve (red line), **B** & **E**. We tested the area between the curves (shaded grey area), the Kolmogorov distance (orange line) and the Euclidean distance between all points on the two curves.

**Figure S12.**
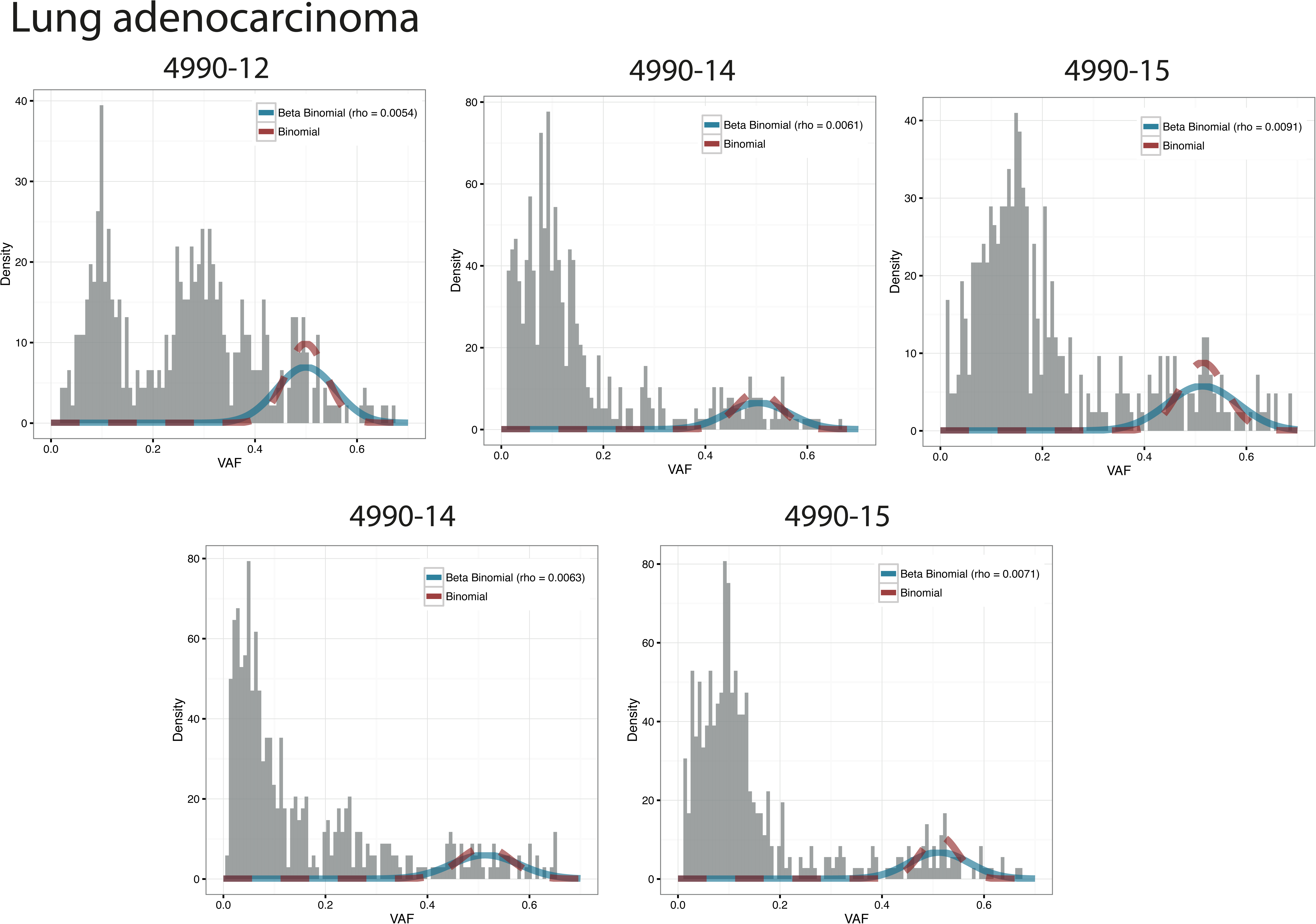
MCMC fits to the clonal clusters of the lung adenocarcinoma samples. We fitted Beta-Binomial and Binomial models to the right hand side of the clonal clusters.

